# Sensing the bactericidal and bacteriostatic antimicrobial mode of action using Raman-Deuterium stable isotope probing (DSIP)

**DOI:** 10.1101/2024.02.12.579891

**Authors:** Jiro Karlo, Arun Sree Vijay, Mahamkali Sri Phaneeswar, Surya Pratap Singh

## Abstract

The mode of actions of antibiotics can be broadly classified as bacteriostatic and bactericidal. The bacteriostatic mode leads to the arrested growth of the cells while the bactericidal mode causes cell death. In this work, we report the applicability of Deuterium stable isotope probing (DSIP) in combination with Raman spectroscopy (Raman DSIP) for discrimination among antibiotics on the basis of their mode of action at community level. We optimized the concentration of deuterium oxide required for metabolic activity monitoring without compromising the microbial growth. We also identified a novel carbon-deuterium Raman metabolic qualitative spectral marker in the biofingerprint region. This can be used for early identification of the antibiotic’s mode of action. Our results explores the new perspective which supports the utility of Deuterium based vibrational tags in the field of clinical spectroscopy. Understanding the antibiotic’s mode of action on bacterial cells in a short and objective manner can significantly enhance the clinical management abilities of infectious diseases and may also help in personalised antimicrobial therapy.

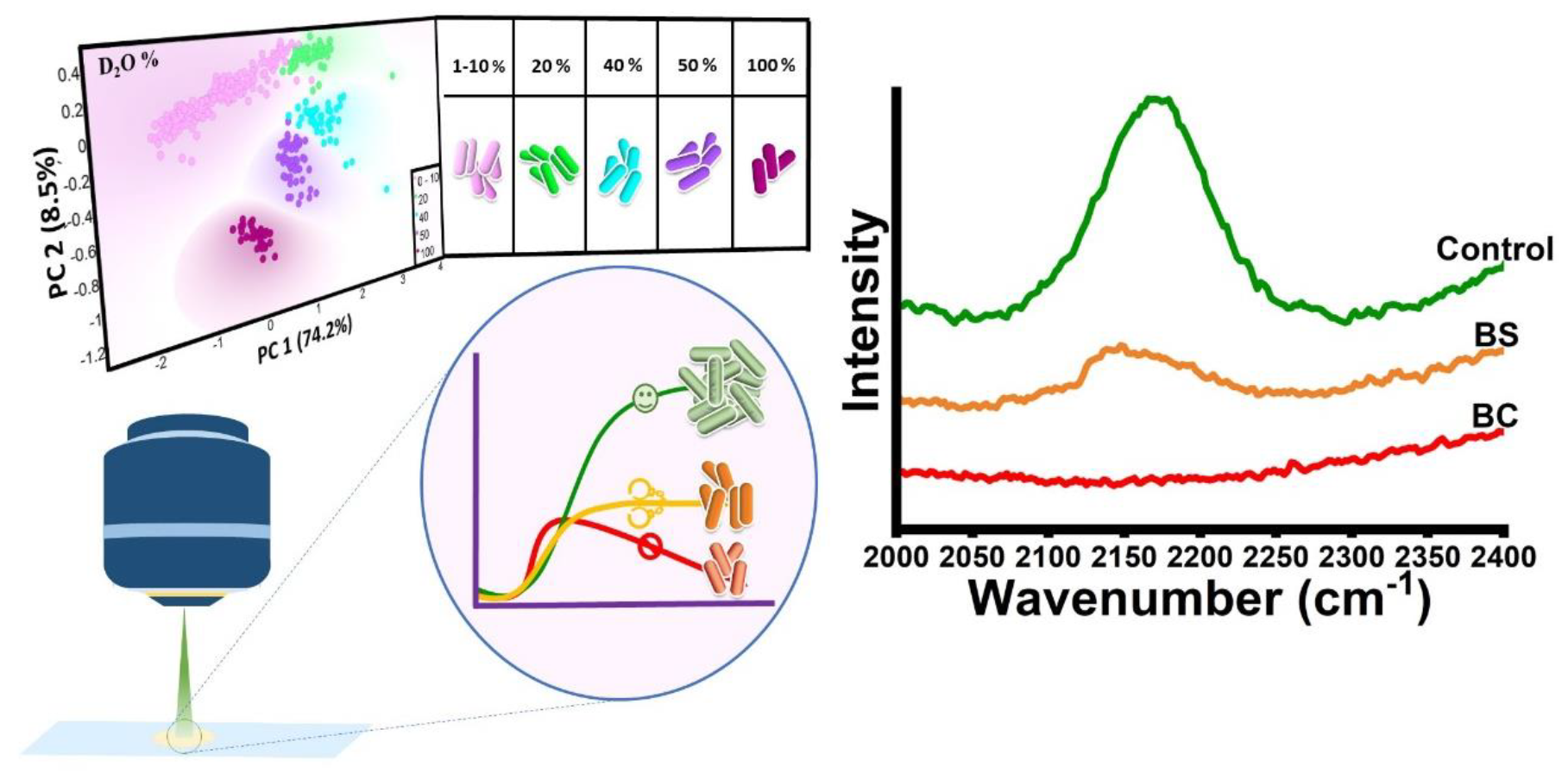

## Introduction

The antibiotic pollution and rampant usage without prescriptions are leading the world to the anti-microbial resistance based major public health issue. ^1–4^ The bactericidal and bacteriostatic efficacy of antibiotics plays an major role in guiding the treatment strategies of bacterial infections. Understanding the differences in the mechanism of action of antibiotics assists in identifying the right antibiotic. Bactericidal antibiotics operates by inducing microbial cell death. It may occur *via* disruption of the cellular structures such as cell walls and other important metabolic functions. In contrast, bacteriostatic antibiotics work by inhibiting cell growth or division without causing immediate cell death. Having said that, bacteriostatic antibiotics also have the ability to kill the bacteria at higher doses.^5^ Commonly used bactericidal antibiotics are fluoroquinolones and β-lactam whereas bacteriostatic antibiotics are chloramphenicol, tetracyclines, trimethoprim/sulfamethoxazole, linezolid, macrolides, clindamycin.^2,6^ Even though the antimicrobial activity of bacteriostatic antibiotics is broad it generally affects the growth of microbes therefore it works synergistically with the host’s immune system to get rid of the microbes.^2,6–8^ While treating the patient with this class of antibiotics, it does not require intensive monitoring with respect to bactericidal antibiotics. ^6,7^ In our study we have used different antibiotics, among them Norfloxacin and Ciprofloxacin are bactericidal (BC) in nature while Chloramphenicol and Tetracycline are bacteriostatic (BS).^2,9^ These antibiotics are used for treatment of many infectious diseases for example; norfloxacin is used to treat acute diarrhea, complicated urinary tract infections (UTIs), gonococcal urethritis; Ciprofloxacin is effective in treating UTIs, gall bladder infections, serious gastroeintestinal infections, eye infections, gonorrohea; Chloramphenicol is effective as conjunctivitis eyedrops, oinment for surgical wounds; Tetracycline is used for treatment of many bacterial infections including acne vulgaris.^10–15^ This broad binary classification system of antibiotics varies in different organisms and the corresponding concentrations^9^. In our study we have used *Escherichia coli* to evaluate the efficacy of these antibiotics. *E*.*coli* are found in the gastrointestinal tract of humans which are rarely infectious in nature unless the host immune system or barriers of the gastrointestinal area are compromised However some of the *E. coli* are pathogenic in nature which causes infections such as diarrhoea, urinary tract infections, meningitis, sepsis.^16^

The existing gold standard method to differentiate between bactericidal or bacteriostatic activity is through optical density measurements.^17^ Another well-known technique is the kill experiment which involves estimating the colony counts on agar plates measuring the microbial cell density^9^. In addition, visual inspection for determining the efficacy of antibiotics is performed using the disk diffusion method and spread plate method. Eventhough, these methods are highly reliable, early detection and biomolecular information cannot be acquired through these techniques. Further just detecting/identifying cells does not necessarily indicate that the cells are metabolically active or viable.^18^ Polymerase chain reaction-based antibiotic susceptibility tests are genotypic methods which are used for faster screening of resistance genes and are culture-free methods but are not suitable for classifying the efficacy of antibiotics on bacterial species^19^.

Deuterium oxide (D_2_O) or heavy water is used as the source of the vibrational probe in the culture medium to substitute the Hydrogen (^1^H) atom with the Deuterium (^2^H/D) in the cellular biomass. When D is substituted in carbon-hydrogen (C-H) bond it gives a unique prominent peak in the Raman silent region which can act as the spectral metabolic activity marker. This also lead to appearance of other unique biomolecular peaks.^20,21^ The appearances of new deuterated peak are the redshifted that arise due to the isotopic effect i.e the substitution by higher mass number D in place of H leading to an increase in the reduced mass of the molecular bond owing to a decrease in wavenumber. ^21–23^ This C-H/D biomolecular substitution occurs due to the rapid intracellular biocatalytic exchange between D of D_2_O and H atom of Nicotinamide Adenine Dinucleotide Phosphate (NADPH) which is redox active.^19,21^ Previously Silvie et al has reported the use of Raman spectroscopy for investigating the action mechanism of bactericidal and bacteriostatic action based on DNA and protein signals in Raman bio fingerprint region (600 -1800 cm^-1^) of *Staphylococcus epidermis*.^2^

In this study, we propose an alternate combinatorial route based on the principles of Raman spectroscopy and Deuterium isotope labelling for identifying the metabolic activity of microbes under the influence of bacteriostatic and bacteriocidal antibiotics. Understanding the mode of action of antibiotics can assist in the clinical management of infectious diseases by customizing the patient’s antimicrobial therapy requirements. This method have the hidden potential for the clinical translation for determining nature of antibiotics and thereby reducing treatment time.

## Methodology

### Bacterial growth and treatment conditions

The bacterial strain *Escherichia coli* K12 was used. It was grown in growth culture medium agar plate incubated at 37°C. A single colony from the culture plate was inoculated in the broth medium and incubated at 37°C with 120 rpm on the rotational shaker. From the overnight culture, re-inoculation was done in the culture medium with different concentration of deuterium oxide concentration – 0%, 1%, 5%, 10%, 20%, 40%, 50% and 100%. Growth profile was determined using OD_600_ value using Epoch 2 microplate spectrophotometer at different incubation time points at 0 hour, 1 hour, 2 hours, 4 hours, 8 hours, 12 hours and 24 hours (2x). Further from the overnight culture, re-inoculation was done in the culture medium with and without antibiotics. The working concentration reported in the previous literatures of the selected antibiotics were used.^2,24,25^ Antiobiotics namely Norfloxacin (Tokyo Chemical Industry), Ciprofloxacin, Tetracycline and Chloramphenicol (Sisco Research Laboratories) were used for the treatment. Growth profile was determined by measuring the OD_600_ value with aforementioned procedure.

### Sample preparation

To determine the optimal D_2_O concentration requirement, aliquots were taken at 18 hours post incoculation from bacterial culture medium with different D_2_O concentrations. The cells were centrifuged at 7000 g for 5 minutes and the pellets were washed twice with phosphate buffer saline (PBS, Sisco Research Laboratories) to remove the media traces. Cell pellets were mounted on ethanol washed CaF_2_ window and air-dried. For the determination of antibiotics mode of action, the washed cell pellets from overnight culture were resuspended in the growth medium supplement with 40% D_2_O with and without antibiotics. Post inoculation the aliquots were collected at different timepoints such as 0 hour, 1 hour, 2 hours, 4 hours, 8 hours, 12 hours and 24 hours. Cell pellets were mounted on ethanol washed CaF_2_ window and air-dried as per the procedure mentioned previously.

### Raman data acquisition and multivariate analysis

For acquiring Raman spectra WITec Confocal Raman Micro spectrometer alpha 300 access was used. It is coupled with a 532 nm laser source, 100x objective with 0.9 numerical aperture and WITec Control 6.1 software. Raman spectra were collected at different spot on the cell pellet and each spectrum was averaged over 10 accumulations with an exposure time of 10 seconds in the spectral range of 200 to 3200 cm^-1^. The acquired spectral data were pre-processed using MATLAB R2021b software. The raw data was smoothen using Savitzky golay filter and baseline corrected with 3^rd^ order polynomial and unit normalized. The Principal component analysis was performed in MATLAB R2021b software using pre-processed spectral data. All spectral plots and figures were generated using Origin Pro 2023b software and the PCA score plot was generate using Orange software.

## Results and Discussion

### Identifying optimal deuteration level for tracking cellular metabolic activity

It has been well established in previously reported studies that the C-D band acts as a prominent Raman metabolic activity spectral marker for cells.^21,26–28^ Determining the optimal D_2_O concentration in the culture medium for the proper growth of the microbe under study without compromising the C-D signal intensity is the most crucial analysis for Raman DSIP-based biosensing. For validating the feasibility of the growth of *E. coli* in D_2_O labelled culture medium, microbial cells were grown in different concentrations starting at 0%, 1%, 5%, 10%, 20%, 40%, 50%, and 100%. Post-inoculation, optical density measurement were performed at different time points; 0 hour, 2 hours, 4 hours, 6 hours 8 hours, 12 hours, and 24 hours. The growth curve was plotted as shown in Figure 1(A). In the growth profile, the low abundance of D_2_O concentration that is 1%, 5%, 10%, 20%, and 40% in the culture medium has a similar pattern of growth curve when compared to growth profile of cells grown in control culture media (no D_2_O). However, with the increase in D_2_O abundance from 50% to 100%, the corresponding growth profile also changes evidently. This retarded growth curved is probably due to the toxicity at higher D_2_O concentrations. Hydrogen (H) bonds play an important role in the biomolecular structure, functions, and interaction. Previously reported studies have reported that substitution of H with D affects the hydrogen bond, disruption of OH/D group orientation and change in symmetry, lipid-protein interactions, protein denaturation process and lipid bilayer.^21,28–30^ At these concentrations, the stationary phase was found to be reaching at earlier time points, suggesting that higher D_2_O abundance has negative affects on the growth and metabolome of the microbial cells.

**Figure 1.**
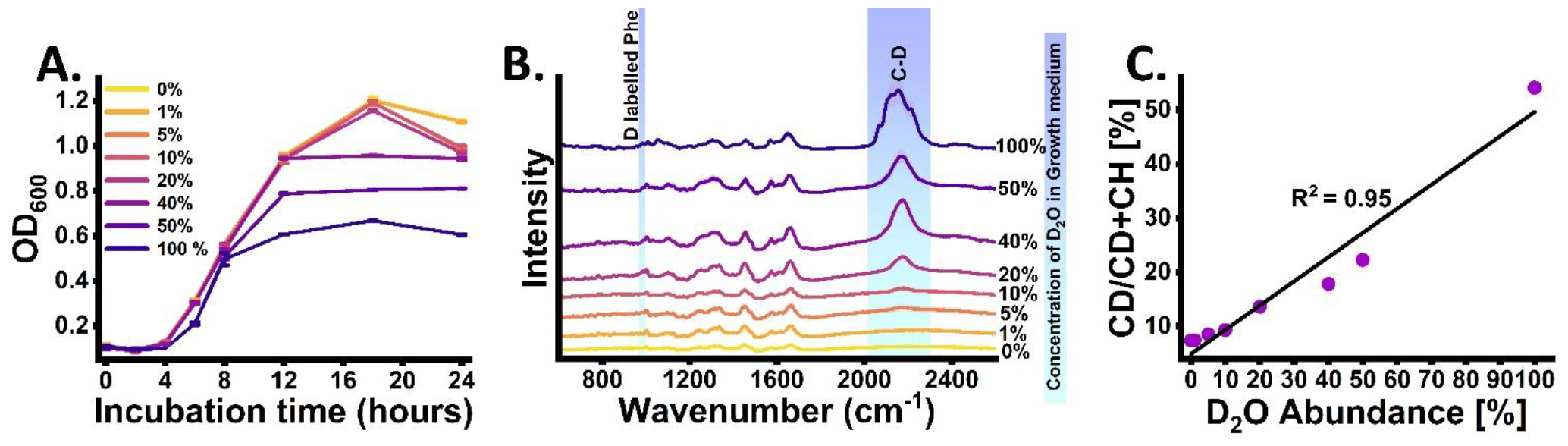
Identifying optimal Deuterium concentration on *E*.*coli* cells; **(A)**. Growth Profile at different concentration of D2O in culture medium with incubation time points. **(B)** Average Raman spectra of *E*.*coli* cells cultured in different D2O abundance **(C)** Plot showing relation between ratiometric CD/CD+CH intensity with D2O abundance in culture medium.

These observations provided evidence that the optimal concentration for microbial growth in presence of D_2_O should be in the range of 1 to 40 %. However, this optimal concentration may vary in different strains. After analysing the growth profile at different D_2_O concentrations, the next step of the analysis was to determine the preferable D_2_O percentage for ensuring a proper detectable signal intensity from the C-D Raman band. Post inoculation, Raman spectra of the microbial cells were recorded at 18 hours time point at different D_2_O percentages as shown in Figure 1(B). In the cells growing with 0 % deuteration, no visible C-D signal was observed in the high wavenumber region between 2070-2300 cm^-1^. Similar trend was observed for cells growing up to 10% D_2_O concentration. A visible C-D peak appeared in cells growing at 20% or higher deuteration level.This new peak arises due to the redshift of the C-H band between 2800-3100 cm^-1^,Table 1. The red shift of the band occurs when protium is substituted by deuterium which has double mass. This increase in reduced mass leads to a decrease in vibrational frequency and the corresponding red shift towards a lower wavenumber ^21,23,26,27^. In Figure 1(B) with the increasing availability of D_2_O abundance 0%, 1%, 5%, 10%, 20%, 40% 50% and 100%, the deuterium-substituted biomolecular Raman bands become prominent. However, the C-D band intensity at 20 % to 100% D_2_O was more prominent when compared to the intensity at D_2_O abundance ranges from 0% to 10%. The observed spectral pattern clearly indicates *de novo* deuterium substitution (H/D) from the culture medium to the cellular biomolecules. Further we have also analysed the biofingerprint region between 600 to 1800 cm^-1^ to analyse the deuteration rate with D_2_O abundance. It has been previously reported that the deuterated phenylalanine, a protein Raman spectral marker have peaks at 989 cm^-1^, respectively. These are redshifted from positions at 1003 and 1126/1168 cm^-1^, respecitvely.^20^ These peaks were visible only in concentrations ranging from 20% to 100% deuteration level. Further the correlation plot between the ratiometric percentage of CD/CD+CH intensity versus the D_2_O abundance shows linear increment relation with 0.95 correlation coefficient R^2^, Figure 1(C).

**Table 1.**
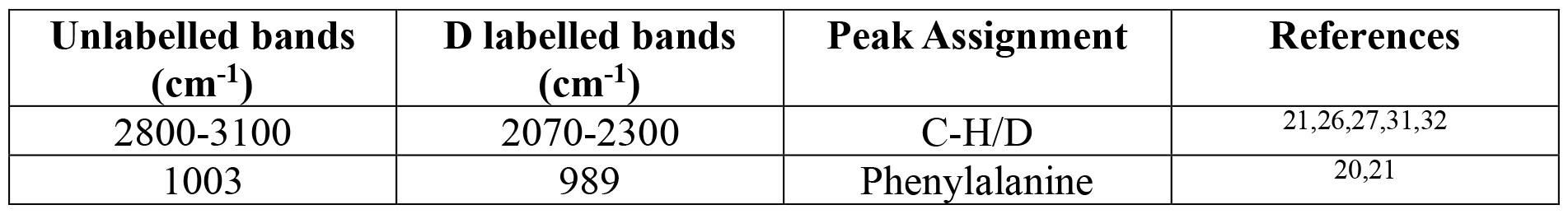
Band Assignments of deuterated biomolecules.

Furthermore to investigate the reproducibility of the Raman spectra from the microbes incubated in increasing D_2_O abundance using the biofingerprint region which have the deuterium substituted biomolecular peaks in much lower intensity when compared to C-D signal in highwavenumber region; we have done a unsupervised statistical analysis using Principal Component Analysis (PCA) as shown in Figure 2. It helps to understand overall variation in the spectral profile. The scatter plot between principal component (PC) 1 and principal component 2 shown in Figure 2 (A) provided distinct clusters for cells treated with higher deuterium concentration. The clusters of 0 to 10% D_2_O abundance were clustered in one group whereas the clusters of 20%, 40%, 50% and 100% clustered with minimal overlapping. The scree plot shown in Figure 2 (B) demonstrated an elbow shaped curve indicating maximum significant variance in the first two principal components which gives 82.7% of the overall variance and the corresponding loading plot is shown in Figure 2(C). Loading plot helps to determine and interpret the biochemical representation by each PC. Comparing the findings of Fig 1(A), 1(B), and 2(A), the 40% D_2_O abundance seems to be the ideal concentration in order to assess a better growth profile and significant C-D band intensity which is subjected to vary from strain to strain.

**Figure 2.**
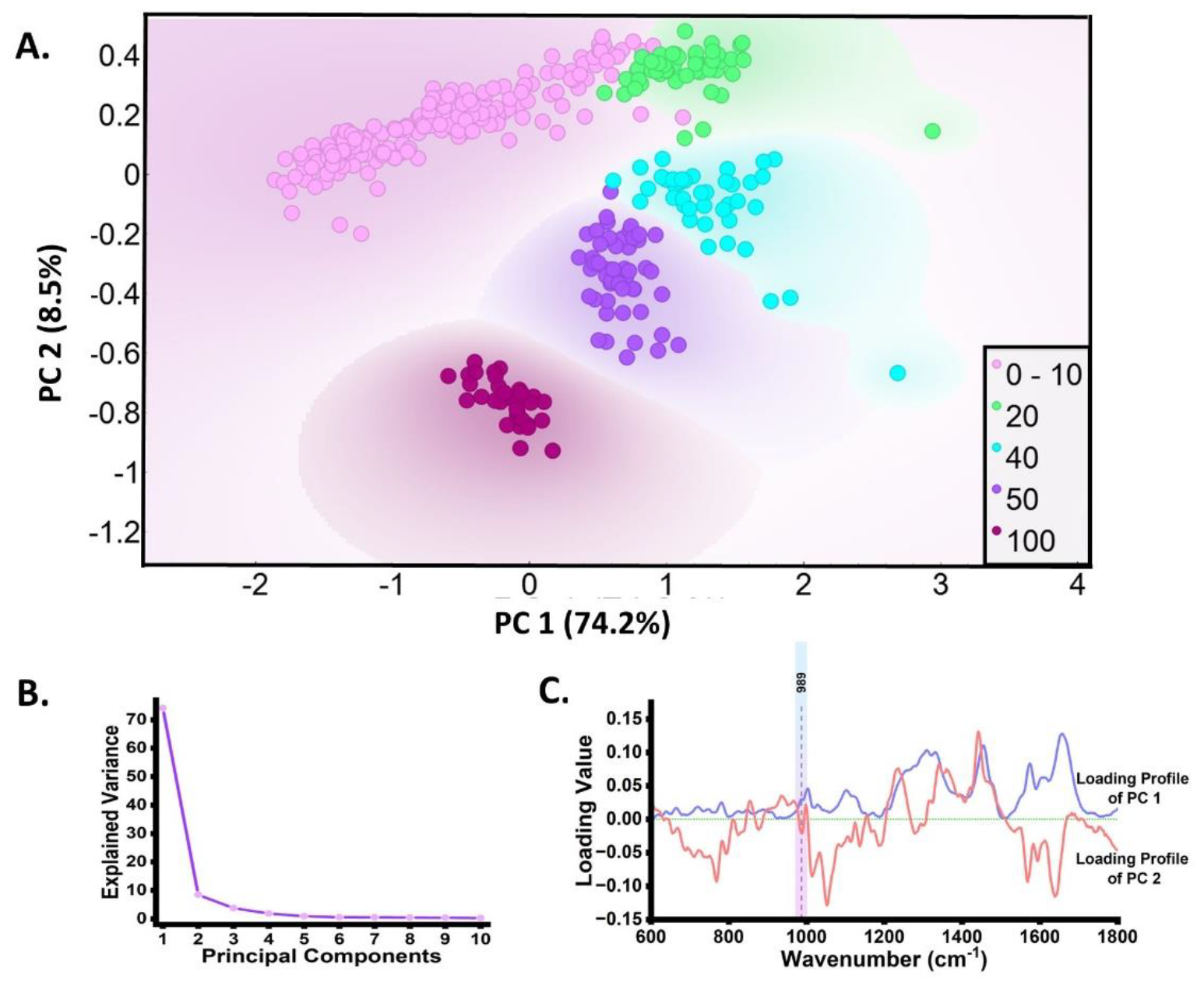
Discrmination of Raman spectra of deuterated *E. coli (biofingerprint region)* cells growing in different D_2_O abundance using Principal component analysis (PCA); **(A)** score plot between principal component 1 and principal component 2; **(B)** Scree plot showing variance contributions by first 10 PCs; **(C)** Loading profile of PC1 and PC2 in the biofingerprint region.

### Bacteriostatic and bactericidal antibiotic activity using Optical Density measurements

To monitor and demonstrate the variability in the antibiotics mode of action, *E. coli* was cultured with and without the antibiotics at 40% D_2_O concentration. The growth kinetics profile through optical density (600 nm) measurement different time points at 0 hour,1 hour, 2 hours, 4 hours, 8 hours, 12 hours and 24 hours was explored, Figure 3A and 3B. Upon monitoring, the untreated microbial growth shows conventional sigmoidal curve. Incubation time of 6 hours was choosen for antibiotic treatment (marked with a star) as microbial culture at this time appear to enter in the early logarithmic phase. In the Figure 3(A), the bacteriostatic antibiotic treated growth curve is displayed where it can be observed that the early stationary phase is reached at 2 hours post antibiotic treatment suggesting inhibited growth of the microbial culture. In contrast, Figure 3(B) where the bacteria has been treated with bactericidal antibioitcs demonstrate halt in the growth and decline of growth curve with incubation time. This declining curve indicates the induction death phase in the culture medium.

**Figure 3.**
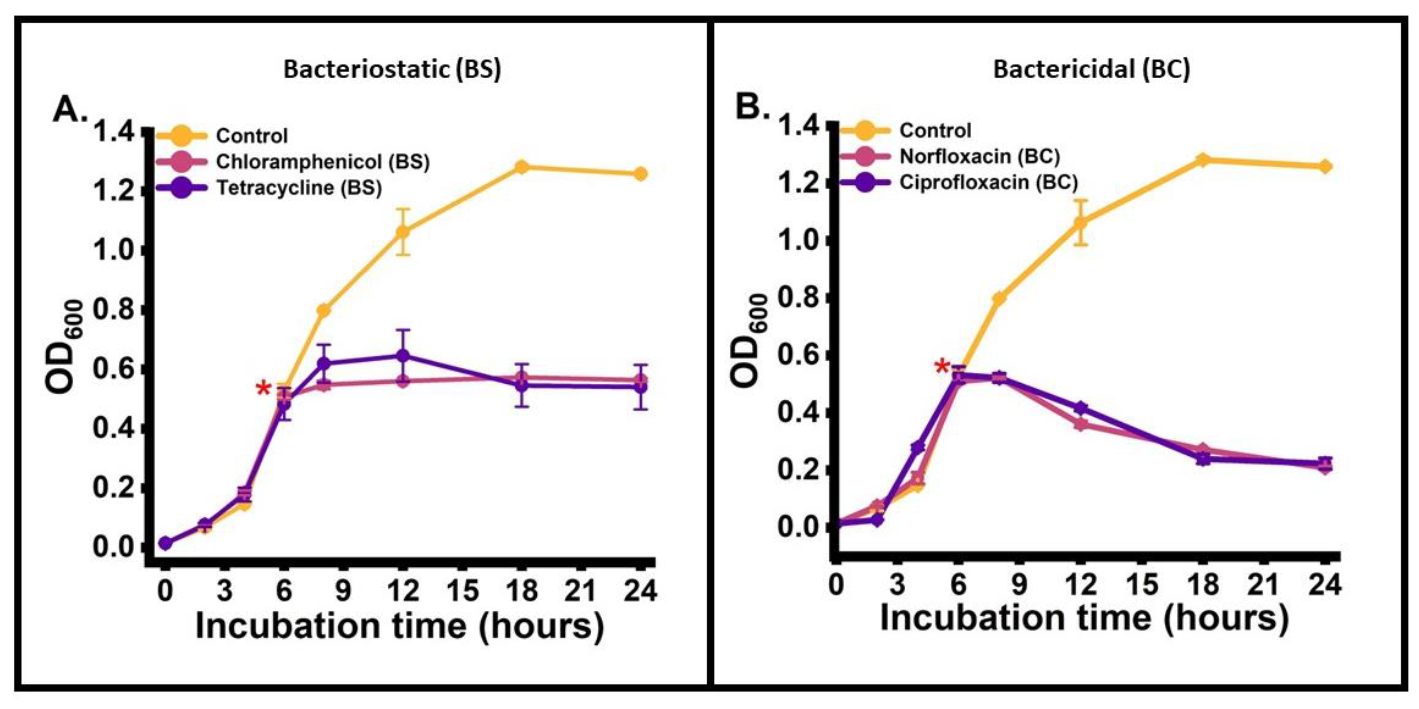
Growth Profile of *E*.*coli* in culture medium with antibiotic treatment at 6 hours**; *(A)*** Bacteriostatic antibiotic (Chloramphenicol and Tetracycline) ***(B)*** Bactericidal antibiotics (Norfloxacin and Ciprofloxacin). [* - Treatment timepoint]

### Raman DSIP in the high wave number region unravels the bacteriostatic and bactericidal antibiotic action

To investigate the action of bacteriostatic and bactericidal antibiotics on the microbial activity using Raman DSIP, the microbial culture medium was prepared with the optimized 40% D_2_O concentration. Before inoculation, the culture medium was treated with different aforementioned antibiotics namely chloramphenicol (30 µg mL^−1^), tetracycline (12.5 µg mL^−1^), norfloxacin (50 µg mL^−1^) and ciprofloxacin (50 µg mL^−1^). Post inoculation Raman spectra of the untreated and treated cells were recorded at different time points starting from 0 hour, 1 hour, 2 hours, 4 hours, 8 hours, 12 hours and 24 hours as shown in Figure 4. At 0 hour, no dectable C-D peak in the region between 2030 to 2300 cm^-1^ was observed for both untreated and treated cells. At the early time points from 1 hour and 2 hours we observe a slight emergence of C-D band in the Raman silent region from the untreated cells. Similarly, we observe emergence of C-D band intensity from the bacteriostatic antibiotic treated cells. In contrast, negligible evergence of C-D band intensity can be seen from the cells treated with bactericidal antibiotic. This qualitative difference in the CD band intensity at early hour indicates the efficacy of this approach to track the differences in the actions of bacteriocidal and bacteriostatic antibiotic. Further, with the increase in the incubation time we observe an intense C-D band from the untreated cells whereas negilibile dynamic changes in the CD band intensity with time is observed for bacteriostatic antibiotic treated cells. At the latter time points, we observed negligible C-D band intensity from the cells treated with bactericidal antibiotic. The negligible C-D band intensity from bactericidal antibiotics treated cells corroborates with cidal action on the cells entering the death phase. For the quantification, we have calculated the band intensity ratio of C-D/C-D+C-H at different incubation time point for the treated and untreated cells as shown in Figure 4(F) and 4(G). The relative intensity difference between control and antibiotic treated cells shows a clear difference with the increase in incubation time points.

**Figure 4.**
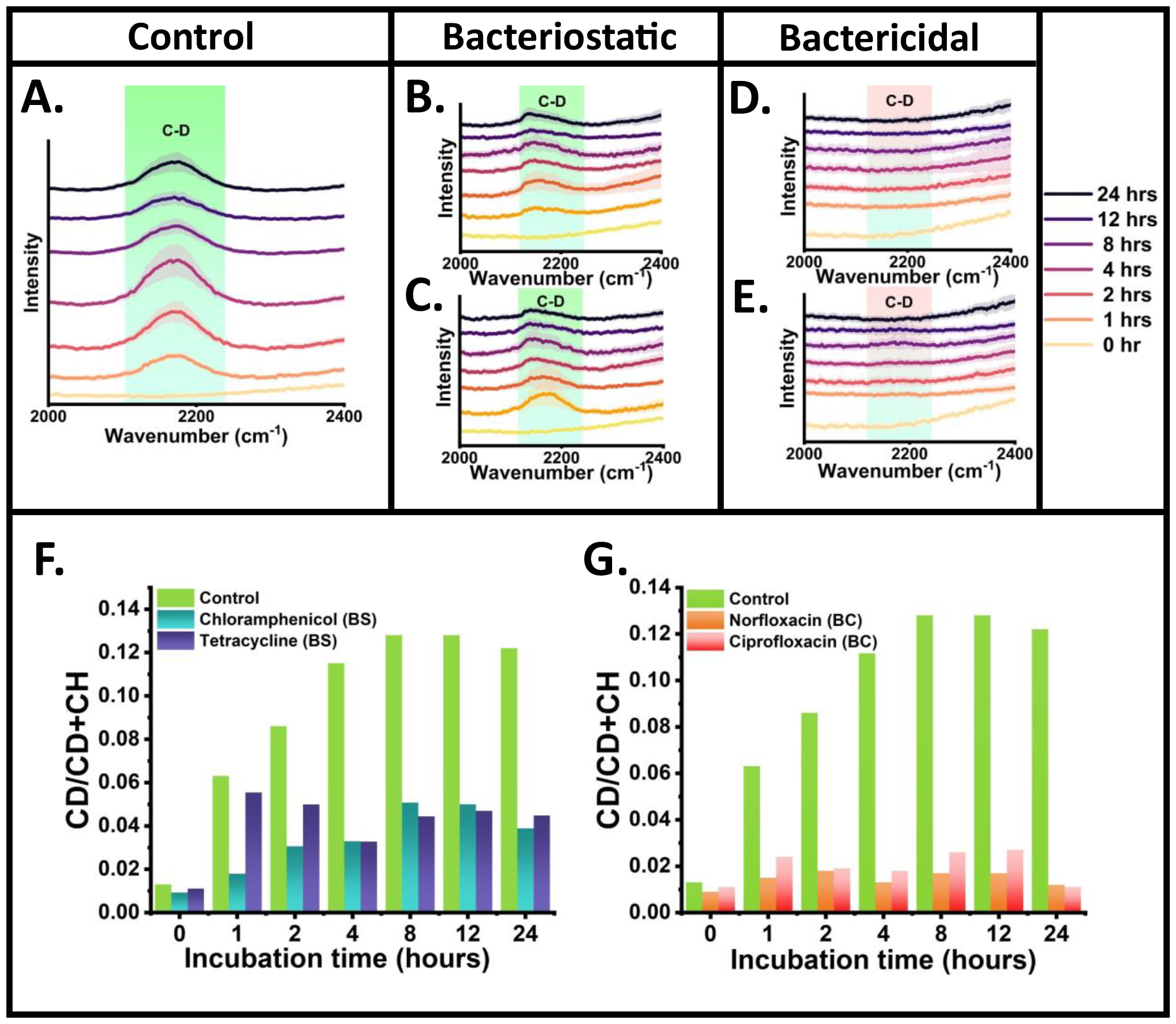
Raman Deuterium Stable Isotope Probing for differentiating the antimicrobial mode of action; Qualitative analysis using Raman spectra of microbial cells at Raman silent reigion in different incubation time points (A) Control (B) Chloramphenicol treated (C) Tetracycline treated (D) Norfloxacin treated (E) Ciprofloxacin treated; Quasi-Quantitative analysis using relative ratiometric intensity (CD/CD+CH) with incubation time (F) Control versus Bacteriostatic treated (G) Control versus Bactericidal treated.

### Raman DSIP in the biofingerprint region to unravel the differences in antibiotic action

Next analysis was to examine whether the D_2_O incorporation also induce spectral changes in the biofingerprint region which can be utilized as metabolic marker to differentiate the antibiotic action. Weak peak observed at 989 cm^-1^ appear due to the deuteration of microbial phenylalanine aromatic ring which is referred a protein reference band.^20,22,28^ Another interpretation of this band at 989 cm^-1^ is to the deuteration of scissoring CH_2_ to leading to the formation of CD_2_.^28^ As the deuterium concentration is taken at 40% complete red shift of this band is not observed. This leads to a visible but weak signal at 989 cm^-1^ which can act as possible qualitative marker band for monitoring deuteration level and corresponding metabolic activity. In Figure 5, in the Raman spectra of control and treated groups at the 989 cm^-1^ position we observe no bands at 0 hours of incubation time. However, this peak appears at 1 hours post incubatiom time point and so on for the control group as shown in Figure 5(A). This indicates the substitution of deuterium in phenylalanine and cellular proteome suggesting towards metabolically active state of cells. In Figure 5(B) Raman spectra for the treated group we observe weak intensity peak at 989 cm^-1^ at 1 hour which becomes weaker with time indicating the bacteriostatic mode of action of action on the cells. Moreover, in Figure 5(C) Raman spectra for the treated group a negligible peak intensity can be seen in all incubation time indicating bactericidal mode of action of antibiotic on the cells.

**Figure 5.**
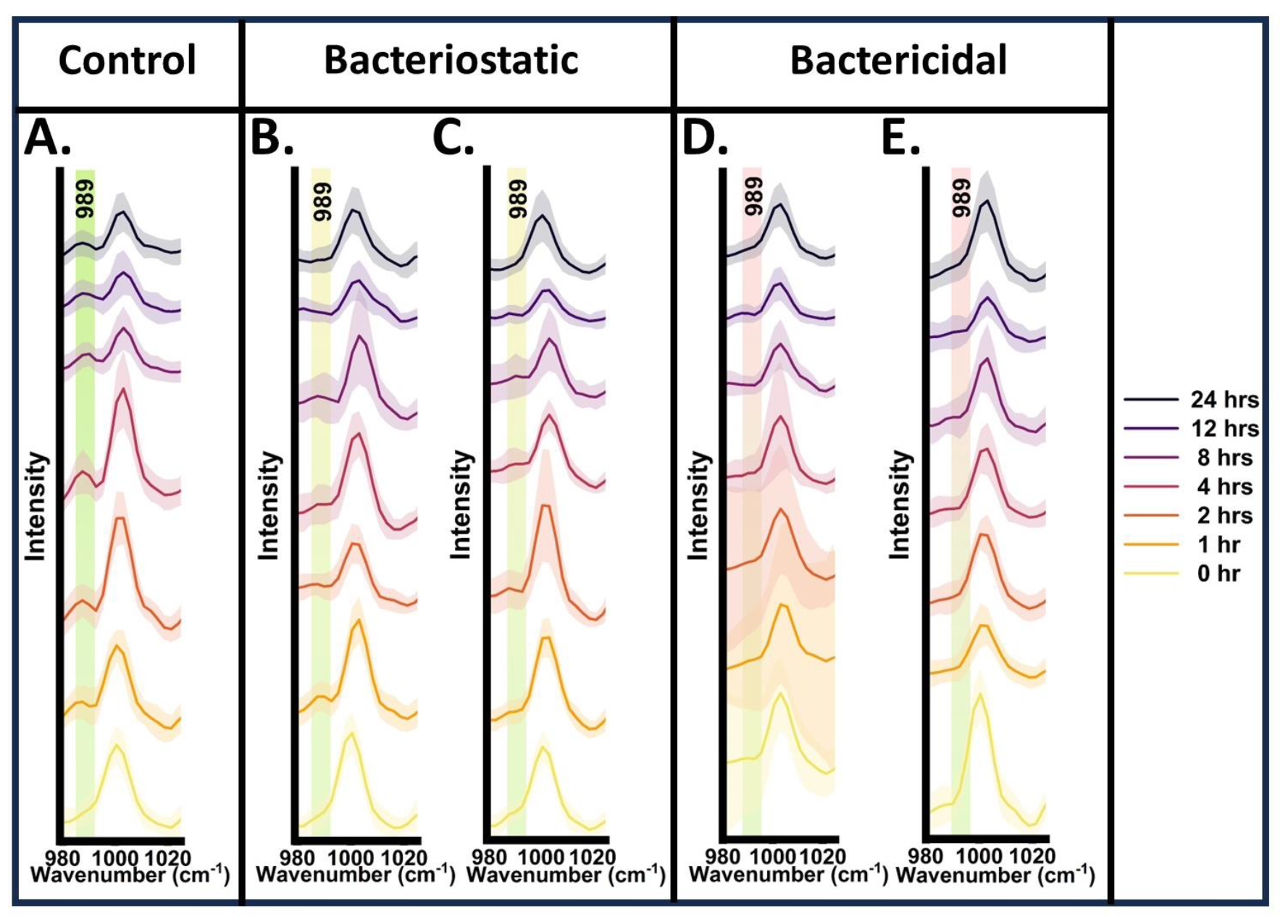
Deuterated phenyalanine band as possible qualitative spectral marker band for differentiating the antimicrobial mode of action; Qualitative analysis using Raman spectra of microbial cells at biofingerprint region at different incubation time points (A) Control (B) Chloramphenicol treated (C) Tetracycline treated (D) Norfloxacin treated (E) Ciprofloxacin treated

Rapid understanding of bacteriostatic and bactericidal antibiotic activity on the microbe can help the clinician to plan appropriate strategies to treat infectious diseases. Antibiotics can be used in accordance to the severity of infection, host immune response, minimizing spontaneous resistant development and potential side effects. In our work, we have used the Raman DSIP as novel assay to rapidly analyse the mode of action of antibiotics. Bactericidal strategy kills the bacteria and bacteriostatic strategy arrest the growth. The obvious difference can be observed in the metabolic activity leading to difference in C-D peak intensity. Raman spectroscopy is a non destructive and non-invasive analytical technique with hidden potentials which require further exploration combining other methodologies such as stable isotope probing.^22,33^ Recently, Raman DSIP based approach has been utilized to detect antimicrobial resistant microbial species. Yang Kai et al has shown the integration of Raman DSIP with advance multivariate analysis to study the spectral dynamics to classify sensitive, intrinsic tolerant, evolved toleratnt and resistant cells.^34^ Zhang Meng et al has reported stimulated Raman DSIP and imaging based strategy to detect metabolic incorporation of D_2_O in single cell level and a rapid antimicrobial susceptibility test (AST) platform in brief span of time of 10 minutes which is claimed to be fastest among currently available technologies.^19^ Xaofie Yi et al has developed a fast Raman-assisted antibiotic susceptibility test (FRAST) based on Raman DSIP for monitoring metabolic activity of bacterial cells (gram-negative and gram-positive) in presence of antibiotics.^35^ David Bauer et al has described a Raman DSIP based AST protocol for gram positive/negative bacteria to detect heteroresistance phenotype and further done proof of concept study by investigating the clinical isolates.^36^ Zhirou xiao et al has reported a AST method based on Raman DSIP to explore efficacy of last resorts antibitotics (tigecycline, polymyxin B and vancomycin) on against *Escherichia coli, Klebsiella pneumoniae, Pseudomonas aeruginosa*, and *Enterococcus faecium*.^37^ Many other recent studies has explored the applicability and potential of Raman DSIP approach for microbial metabolic activity and antimicrobial studies.^38–41^ Although multiple studies have utilized D_2_O concentration from 40 to 70%, we first optimised the feasible concentration which is necessary for best survival of microbe keeping a detectable C-D band under consideration. One of the remarkable findings of this study was to analyse the deuteration effect in the biofingerprint region and apply the multivariate analysis using PCA with classified clusters showing difference in percentage of deuteration in cells. Further the bacteriostatic and bactericidal effect was observed using spontaneous Raman scattering at a very early incubation time point. Though Raman DSIP is a highly feasible non destructive approach the D_2_O has toxic effect on the cells which we have shown and is the limitation in our study. Future perspective of Raman DSIP approach lies on tracing the metabolic activity by increasing C-D band detection sensitivity at very lower concentrations of D_2_O and in a short span of incubation time. This will minimize the toxic effect on the microbial metabolome and will promote Raman DSIP as a cost effective approach to develop into a translational clinical assay for antimicrobial studies.

## Conclusion

Our results suggests the dynamic changes in the intensity of Carbon-Deuterium band in Raman silent zone of high wavenumber region can differentiate between the metabolic active, growth arrested and cidal microbes. Further, we explored the possible potential of deuterated biomolecules band in the biofingerprint region as metabolic spectral marker for qualitative analysis of our mentioned goal. This work project Raman DSIP as a prospective alternate biosensing assay to determine the efficacy of antibiotic on the microbes at the community level. With this approach, the effect of antibiotics can be determined at early incubation time points. As the proposed method is efficient and simple, it hold immense potentials for the clinical translation as a tool for rapid classification of the antibiotics and concentration dependent effect as bactericidal or bacteriostatic. The exploration of Raman DSIP holds the potential to be translated into a rapid clinical assay for selection of appropriate antibiotics to treat infectious diseases, detection of antimicrobial resistance and microbial dormancy.

## Acknowledgements

This work was carried out under research grant project no (37/1739/23/EMR-II) supported by Council of Scientific and Industrial Research (CSIR), Government of India and project no. IIRP-2023-1734 from Indian Council of Medical Research (ICMR), Government of India.

